# Spatial Perspective Taking is Distinct from Cognitive and Affective Perspective Taking

**DOI:** 10.1101/2024.06.16.599248

**Authors:** Maria Brucato, Nora S. Newcombe, Jason Chein

## Abstract

Perspective taking (PT) is the ability to imagine viewpoints different from our own. However, the nature of PT as a construct, and its underlying cognitive mechanisms, are not well established. Some researchers propose that understanding what others believe (cognitive PT), feel (affective PT), and see (spatial PT) form a single behavioral dimension, relying on the orienting of attention between competing frame-of-reference representations. Others propose that PT mechanisms are dissociable, although there are three different proposals about such dissociations. The present study examined behavioral associations among measures of spatial, cognitive, and affective PT and attentional control in neurotypical young adults. There was a lack of convergent validity for measures of cognitive and affective PT, pointing to the need for more psychometric work on these dimensions. Much better convergence was found for spatial PT measures. There was little to no behavioral association between spatial PT and either social forms of PT (cognitive or affective) or attentional control measures. This pattern suggests support for a dissociated model in which spatial PT is a distinct cognitive construct.

## Spatial Perspective Taking is Distinct from Cognitive and Affective Perspective Taking

Perspective taking (PT) is the name often given to a psychological construct that includes the ability to imagine another person’s experience in one of several ways: what others see (spatial PT), what they believe (cognitive PT), and what they feel (affective PT). Spatial PT supports successful spatial navigation of large-scale environments (Allen et al., 1996; Fields & Shelton, 2006; Kozhevnikov et al., 2006; Nazareth et al., 2018) and may also contribute to educational achievement and career success in science, technology, engineering, and mathematics (STEM) disciplines (Oldakowski, 2001; Plummer et al., 2016; Wai et al., 2009). Cognitive and affective PT are important for forming and maintaining high-quality social relationships (Galinsky et al., 2005; Hughes & Leekam, 2004; Long & Andrews, 1990). Deficits in cognitive and affective PT are also characteristic of many clinical populations (Cotter et al., 2018), including autism spectrum disorder (Baron-Cohen, 2000) and major depressive disorder (Bora & Berk, 2016), and may play a role in development, progression, and responses to treatment (Bora & Zorlu, 2017; Quednow, 2017). Differences in the severity of PT impairments may be related to general mechanisms that support this ability regardless of diagnostic category and may help connect and differentiate between traditional diagnostic categories as in a Research Domain Criteria approach (Cuthbert & Insel, 2013). However, it is currently debated whether PT constitutes a singular psychological construct, and if so, what cognitive mechanisms support it.

### Does Perspective Taking Constitute a Singular Psychological Construct?

There are four views on the question of whether spatial, cognitive, and affective PT cohere to form a single dimension (see Figure 1). The first view, the “distinct” account, proposes distinct mechanisms for all three kinds of PT. For example, modularity theories regarding theory of mind claim that there is an innate domain-specific neural mechanism dedicated to social mental state reasoning (e.g., Carruthers, 2013; Leslie et al., 2004). In line with this theory, some researchers hypothesize that the temporoparietal junction (TPJ) is associated with reasoning about beliefs (cognitive PT) but does not support other cognitive tasks (e.g., spatial PT; Saxe & Kanwisher, 2004; Saxe & Wexler, 2005; Kosakowski & Saxe, 2018). A second proposal, the “spatial vs social” account, is drawn in part from studies of individuals with autism spectrum disorder and aligns with Baron-Cohen’s (2009) empathizing-systemizing theory. In this view, cognitive and affective PT (social forms of PT) have distinct cognitive mechanisms from spatial PT. In support of this account, a recent neuroimaging meta-analysis found that cognitive and affective PT showed large regions of overlap in functional activation, whereas both showed virtually no overlap with spatial PT (Brucato, 2022). A third proposal, the “cold vs hot” account, suggests that distinct mechanisms may support PT tasks that require representation of “cold” versus “hot” mental states (Aichhorn et al., 2006). In this view, spatial and cognitive PT are purported to involve reasoning about non-emotional or “cold” representations, commonly engaging the posterior end of the superior temporal sulcus and TPJ. Conversely, affective PT, and to a lesser extent cognitive PT, which both require individuals to represent and predict emotional or “hot” mental states, are thought to rely additionally on neural mechanisms in the medial prefrontal cortex.

**Figure 1.**
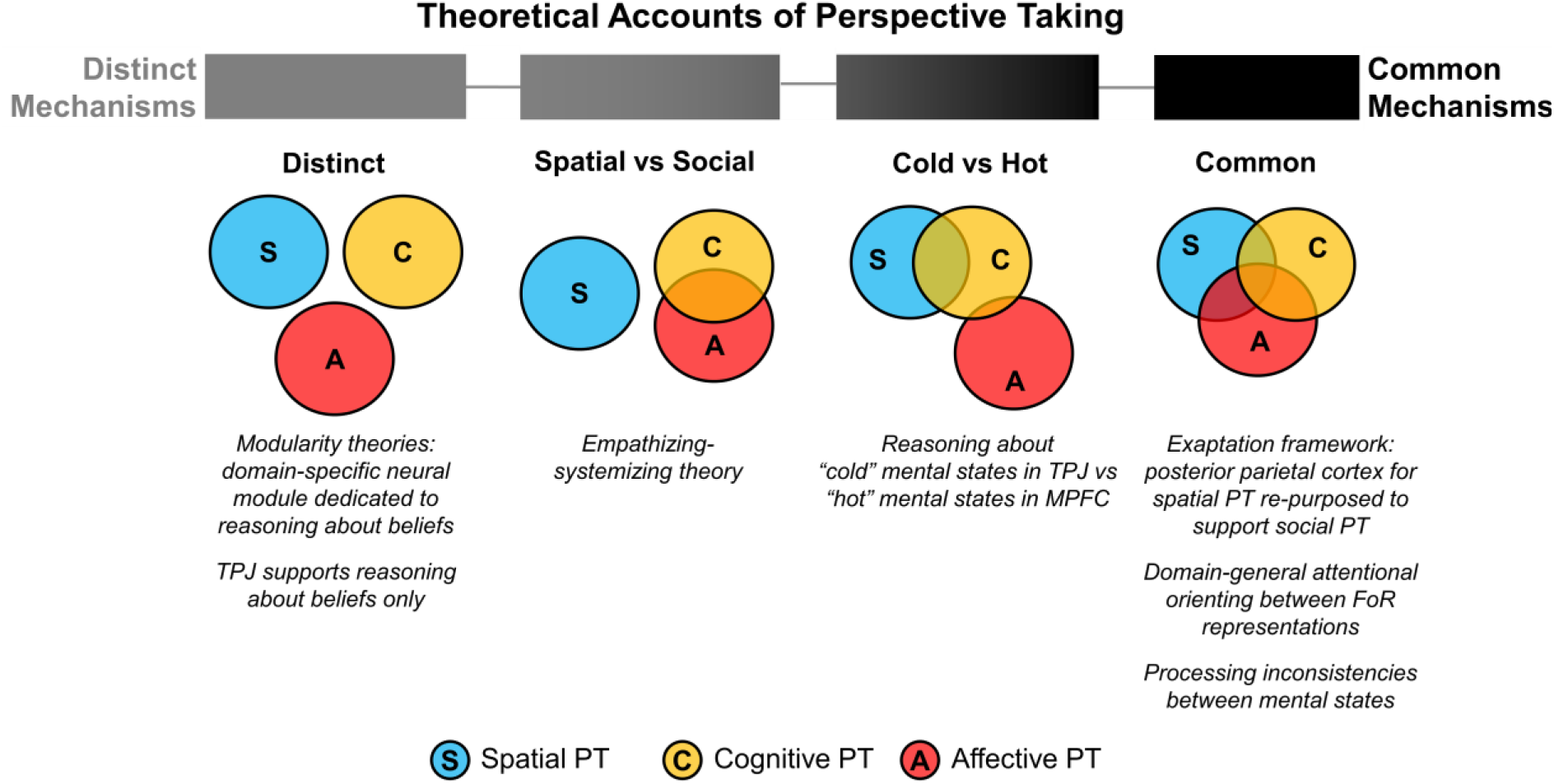
Schematic Overview of Theoretical Accounts of Perspective Taking. *Note.* PT = perspective taking; FoR = frame-of-reference; TPJ = temporo-parietal junction; MPFC = medial prefrontal cortex. The top bar gradient depicts how closely each account aligns with distinct mechanisms accounts (lighter) or common mechanisms accounts (darker).

A competing set of theoretical and empirical papers propose a fourth, unitary, or “common” account in which spatial, cognitive, and affective PT all rely on a common set of cognitive processes (e.g., Arora et al., 2017; Parkinson & Wheatley, 2013; Quesque & Rossetti, 2020; Sun & Wang, 2014), albeit for varying reasons. Parkinson and Wheatley (2013), for instance, argued that evolutionarily older neural circuits used for reasoning about physical space were later re-purposed for social cognition. Others postulate that spatial, cognitive, and affective PT are unified because they all require processing inconsistencies between mental states (e.g., Arora et al., 2017; Quesque & Rossetti, 2020; Schurz et al., 2013) or because each involves the manipulation of frame-of-reference representations, which leads to shared reliance on dorsal and ventral attentional networks (Sun and Wang, 2014). That is, attention orienting could be a domain-general mechanism that supports the comparison or shifting between alternative possible perspectives, regardless of domain.

With clinical populations, research shows a compelling pattern of co-deficits among cognitive, affective, and spatial PT, favoring the unitary view. Autism spectrum disorder is characterized by difficulties in understanding the mental states of others, on measures of cognitive (Baron-Cohen, 2000; Yirmiya et al., 1998), affective (Song et al., 2019), and spatial PT (Hamilton et al., 2009). Further, cognitive PT has been found to significantly predict spatial PT performance for both autism spectrum disorder and typically developing children (Hamilton et al., 2009). Individuals with major depressive disorder are also known to have impairments in cognitive and affective PT, and greater symptom severity is associated with worsening cognitive and affective performance (Bora & Berk, 2016). Further evidence suggests that individuals with sub-clinical depression may also have co-deficits in spatial PT (Erle et al., 2019).

The bulk of the research on PT dimensionality, however, involves neurotypical adults, and the evidence here is much more mixed (see Table A1 & A2). Four early experiments assessed the association of self-reported affective PT and reaction time on a spatial PT task called the Own Body Transformation Test, and found differing results (Gronholm et al., 2012; Mohr et al., 2010; Thakkar et al., 2009; Thakkar & Park, 2010). Two of these experiments found a significant correlation between self-reported affective PT and reaction time on the spatial PT task for female but not male participants; however, the effect was directionally opposite between the two studies (Mohr et al., 2010; Thakkar et al., 2009). Further, neither study found an association between self-reported affective PT and spatial PT accuracy (Mohr et al., 2010; Thakkar et al., 2009). Thakkar and colleagues (2010) later reported that more spatial PT errors were correlated with lower self-reported affective PT in female participants but failed to replicate reaction time results of their earlier study. Across four experiments, Erle and Topolinski (2015) found that greater spatial PT accuracy and quicker reaction time was significantly associated with higher self-reported affective PT in neurotypical adults. However, none of these studies measured objective affective PT performance. Meanwhile, relatively fewer studies have investigated the association of cognitive PT and spatial PT (Table A2). Two recent studies used objective measures of these two types of PT and found contrasting results either indicating a significant association (Putko & Złotogórska -Suwińska, 2019) or not (Job, 2021).

Importantly, most studies have not tested all forms of PT together in one sample. The fact that all three types of PT are not tested together limits our ability to use this evidence in distinguishing between common and distinct mechanisms accounts of PT. Indeed, extant behavioral research on the question of common or distinct dimensions of PT, and the role of attention orienting in supporting PT across different domains, renders a murky picture. Broadly, work in adults produces inconsistent findings, sometimes pointing to shared processing, and sometimes not.

Work conducted in children also fails to clarify the situation. One early review on children’s PT abilities concluded that performance on PT tasks exhibits little to no between-domain correlations (Ford, 1979). However, more recent work has pointed to behavioral covariance across PT domains, including evidence for an association between cognitive and spatial PT in very young children (Hamilton et al., 2009; Tian et al., 2021; Viana et al., 2016), and evidence of significant associations between cognitive and affective PT in both younger (Bensalah et al., 2016), and older (Cassetta et al., 2018) children. Aslan and Köksal Akyol (2020) developed a PT measure for children designed to assess all three types of PT abilities in one test. The fact that the overall measure exhibited good internal consistency (α = 0.71) suggests that spatial, cognitive, and affective PT items in the test must have covaried substantially in the sample (see also Tian et al., 2021), and thus, offers some support for a unified view of these abilities.

### Convergent Validity Within Perspective-Taking Domains

Many different assessments exist for each PT construct, and each has an unclear and largely untested relations to others, even within a given PT domain. Few studies have administered more than one task of the same type of PT to establish convergent validity either within or between PT domains, which is crucial when considering previously reported issues of within-domain convergent validity for social PT tasks (Ford, 1979; Gallant et al., 2020; Quesque & Rossetti, 2020; Warnell & Redcay, 2019). For example, prior reports investigating the construct validity of Theory of Mind tasks have found that there is none (i.e., Warnell & Redcay, 2019, Gallant et al., 2020). Theory of Mind is a psychological construct defined as the ability to understand others’ emotional or cognitive mental states and differentiate them from one’s own (i.e., social PT). However, it has recently been argued that some of the most commonly used Theory of Mind tasks do not require PT at all (Quesque & Rossetti, 2020). Inclusion of tasks which do not meet criterion as PT tasks may explain previous absences of convergent validity within PT domains.

Navarro (2021) conducted the first psychometric investigation of Theory of Mind tasks, including the Director Task, the Short Stories Questionnaire, and the Reading the Eyes in the Mind Task. There were small significant correlations among all tasks in the battery, but psychometric network analysis indicated that these Theory of Mind tasks did not represent a unified cluster. Specifically, the Reading the Eyes in the Mind task did not share any significant edges with the Director Task. In some ways this finding is unsurprising, since other work has concluded that the Reading the Eyes in the Mind Task is actually a measure of emotion recognition and not explicitly a PT task, insofar as it does not require representation of mental states or differentiation between one’s own and another’s perspective (e.g., Oakley et al., 2016; Quesque & Rossetti, 2020).

Intentional investigations into the psychometric properties and convergent validity of popular affective PT tasks in neurotypical adults are sparse. The Yoni Task (Shamay-Tsoory et al., 2007) has recently been shown to demonstrate strong psychometric properties, and affective PT trials of the Yoni Task significantly correlated with the Reading in the Minds Eyes Task (r = 0.29), but not with a gender discrimination task (Isernia, Rossetto, Blasi, et al., 2023; Isernia, Rossetto, Shamay-Tsoory, et al., 2023). Given the claim that Reading in the Minds Eyes Task may not require PT (Oakley et al., 2016; Quesque & Rossetti, 2020), though these findings are promising with respect to the validity of social skills measures, they do not necessarily provide information about the convergent validity of the Yoni Task with other measures of affective PT.

Meanwhile, in the spatial domain, PT is inconsistently tested as what Flavell (1981) called Level 1 spatial PT (establishing a line of sight from another’s eyes) and Level 2 spatial PT (working out the details of what another sees). Promisingly, a recent psychometric analysis of behavioral tasks used to measure Level 2 spatial PT reported that at least three of four commonly used behavioral measures of this construct show strong psychometric properties, including good item discriminability and internal reliability within-tasks, convergent validity between tasks, and divergent validity from other spatial skills like object rotation (Brucato et al., 2022).

In addition, for all three kinds of PT, there are both objective assessments and self-report measures, but the extent to which self-reported PT abilities reflect performance on objective tests is variable (Baksh et al., 2018; Ickes, 1993; Israelashvili et al., 2019; Murphy & Lilienfeld, 2019; Murray et al., 2017; Sunahara et al., 2022).

### Attentional Control and PT Performance

Despite assertions from researchers working within spatial, cognitive, and affective domains about the potential role of attentional control in supporting PT performance (Amorim et al., 2003; Wade et al., 2018), there are very few studies that directly test these claims. A small collection of behavioral studies conducted with adults and children have found positive associations between attention switching and Level 2 spatial PT (De Lillo & Ferguson, 2023), cognitive PT (Austin et al., 2014; Long et al., 2018) and affective PT abilities (Goodhew & Edwards, 2021). However, little is known about whether behavioral measures of attentional control might play an explanatory role in the occasionally observed behavioral co-variance of spatial, cognitive, or affective PT abilities. The present study attempts to bridge these gaps in behavioral work to more adequately adjudicate between common and distinct mechanisms accounts of PT.

## Method

### Participants

Participants were 221 undergraduates at Temple University and adults living in the surrounding area (139 female; 71 male; 10 non-binary; 1 did not report gender). The mean sample age was 19.48 (SD = 3.22 years), and ranged from 17 to 41 years. All participants had normal or corrected-to-normal vision and hearing. List-wise deletion was used to address missing data. Six participants had missing data on one task or questionnaire in the behavioral battery and were removed from the dataset. Thirteen participants failed comprehension checks for at least one task and/or two or more attention checks in questionnaires and were thus also removed from the dataset. Univariate and multivariate outlier analyses were also conducted. Specifically, seven participants were identified as univariate outliers as they had a mean score on at least one task that was three or greater standard deviations from the sample mean. Two multivariate outliers were also identified based on extreme Mahalanobis distance (p < .001 for the χ^2^ value). In total, nine outliers were removed from the dataset. The final sample contained 193 participants (120 female; 63 male; 10 non-binary) with a mean age of 19.44 years (SD = 3.07 years). All participants gave informed consent before completing the protocol, which was approved by Temple University’s Institutional Review Board. As compensation, participants received either partial course credit or a $30 gift card following the study session.

### Procedure

Over a study session that lasted approximately two hours, participants completed six PT tasks, an attention task, a reasoning task, and three self-report measures. After providing informed consent, participants were randomly assigned to one of 10 possible testing orders that counterbalanced tasks to avoid confounding due to task sequencing or fatigue effects. Each testing order was arranged such that PT tasks of the same domain did not directly follow one another, and each task was first and last in the behavioral testing battery exactly once. All measures were administered on a Windows 64-bit computer with an LCD monitor display. Up to two participants were tested at once in the testing room. Testing computers were on opposite sides of the same wall of the testing room with a separator in between, and participants wore headphones during the experiment so that any tasks that contained an audio stimulus did not disrupt the testing session for either participant.

### Materials

#### Perspective Taking Tasks

Two behavioral tasks were selected to measure individual differences in perspective taking in each domain. In the spatial domain, one task (Perspective Taking Task for Adults) contained stimuli that resembled a human (i.e., an "agent”) and the other did not (Spatial Orientation Task). These tasks were selected under the assumption that spatial PT tasks including a “social” agent might co-vary more substantially with social PT tasks (see Shelton et al., 2012; Clements-Stephens et al., 2013). Applying the same logic, one cognitive PT (Director Task) and one affective PT (Yoni Task) task were selected because they contained stimuli with more spatial features, and the alternative cognitive (Strange Stories Task) and affective PT (Empathic Accuracy Task) tasks were selected to have stimuli with fewer spatial features.

##### Spatial Orientation Task

The Spatial Orientation Task (Friedman et al., 2020) was used as a measure of spatial PT. The task displayed a configuration of two-dimensional objects on the left side of the screen and a response circle on the right. At the start of each trial, participants were asked to imagine that they were standing at one object and facing a second object. Their imagined position was represented by a dot in the center of a response circle, and their imagined line of sight by a solid arrow pointing upwards. Participants were then asked to imagine pointing towards a third object and to drag the arrow in the response circle to indicate their imagined pointing direction. Participants completed four practice items followed by 12 test items in a random order. The angular error between the correct response and the participant’s response was measured for each trial. Participants were allotted 5 minutes to complete the task. Participants received an angular error score of 90 degrees for each item that was not answered in the allotted time. The total score was equal to the average angular error across all trials.

##### Perspective Taking Task for Adults

The Perspective Taking Task for Adults (Brucato et al., 2022) was used as another measure of spatial PT. Participants saw an image of a three-dimensional human-like figurine taking a photograph of an arrangement of three different-colored three-dimensional objects. Eight different perspectives of the arrangement were simultaneously provided below this image, and participants were asked to determine which one looked like the picture the figurine could have taken from where it was standing. There were three practice items followed by 32 test items, presented in a fixed quasi-random order. Participants were allotted 3 minutes to complete all test items. Four additional items, in which the perspectives of the figurine and participant were the same (first-person-perspective,1PP) were used as comprehension checks for attentive responding, such that incorrect responses to two or more 1PP items resulted in exclusion from the analysis. The total score for the test was the number of correctly answered items per minute, excluding 1PP items.

##### Empathic Accuracy Task

The Empathic Accuracy Task (Mackes et al., 2018) was used as a measure of affective PT. Videos of individuals (targets) recounting emotional autobiographical events and those targets’ emotional intensity ratings during each recollection were acquired from the Free Emotional Event Library (Mackes et al., 2018). Six video clips were presented to each participant: 2 happy, 2 sad, and 2 angry. One video of each emotion had a male target and the other had a female target. Video clip presentation was counterbalanced using a balanced Latin square design. One of the six counterbalanced video clip orders was randomly assigned to each participant. While watching each clip, participants were asked to continuously rate the emotional intensity of the target on a 9-point scale (from 1, ‘no emotion’ to 9, ‘very strong emotion’) using the keyboard. After each clip finished, participants were asked to select (1) which emotion the target felt most strongly (options of “happy”, “angry”, “surprised”, “sad”, “frightened” and “no emotion”), and (2) which emotion they themselves felt most strongly. The analysis outlined in Mackes et al., (2018) was followed to derive empathic accuracy scores for each participant. First, ratings for each video were separated into 2-second bins and one time-weighted average rating was calculated for each bin. Next, Pearson’s correlation coefficients were calculated for the association between participants’ and targets’ ratings. The resulting Pearson’s correlation coefficient for each video clip and each participant was then r-to-Z transformed to allow comparison between correlation coefficients.

##### Yoni Task

The Yoni Task (Shamay-Tsoory et al., 2007) was also used to measure affective PT. There were 98 test trials that showed a cartoon outline of a face (named “Yoni”) surrounded by four pictures in each corner of the computer screen of objects belonging to a single category (e.g., fruits, chairs) or faces. Participants were asked to click on the correct image to which Yoni is referring as quickly and accurately as possible. The stimuli contained two types of cues that participants could use to make their responses: verbal only (i.e., a sentence that appeared at the top of the screen) or verbal cue with eye gaze. On each trial, Yoni’s eyes looked at the correct answer (55 trials), straight ahead (32 trials), or at a misleading answer (11 trials). There were 3 main trial types: cognitive, affective, and physical. The cognitive and the affective trials involved mental state inferences, whereas physical trials required a choice based on a physical attribute of the character. In addition, each trial type had two levels of difficulty: first order and second order. In first order trials, participants were only required to understand Yoni’s mental state (e.g., “Yoni loves ”), whereas second order trials required understanding the interaction between Yoni’s mental state and each of the four characters around him (e.g., “Yoni loves the animal that loves”). Participants who scored less than 50% on the physical control trials were removed from analyses involving this task. Accuracy of affective trials was summed to get an affective PT score ranging from 0 to 48.

##### Strange Stories Task

The revised Strange Stories task (White et al., 2009) was used to measure cognitive PT. During each trial, a short story or a series of unlinked sentences appeared on the screen. After reading the text, participants pressed the space bar to receive a question. On mental state trials, participants were asked a question that required them to reason about the mental state of characters in a short story, and on unlinked sentences trials participants were asked to report a fact from one of the unlinked sentences. There were 8 mental state trials and 8 unlinked sentences trials, presented in a randomized order. Participants were instructed to wait until they thought of the correct answer and then to press the space bar to advance. They could then type their response into a text box on the screen and hit the “Enter” key to advance to the next trial. Reaction time was recorded during the think period for each trial. Typed responses to each answer were then coded for accuracy by two independent raters, on a scale of 0 to 2 based on the scoring paradigm from White et al., (2009). Following White and colleagues’ protocol, 20% of the scores selected randomly for each story were used in an intraclass correlation analysis to determine level of agreement between raters, and good agreement was reached (intraclass correlation coefficient = .83). Participants who scored less than 50% on the unlinked sentences stories were removed from analyses involving this task. Accuracy on mental state story trials were summed for a cognitive PT score ranging from 0 to 16.

##### Director Task

The Director Task (Dumontheil et al., 2010) was also used to measure cognitive PT. The stimuli showed shelves (4 slots high and 4 slots wide) with a director standing behind them. Eight objects (some were different types of the same kind of object: e.g., golf ball, tennis ball, basketball) were shown in slots of the shelves, and 5 slots were occluded such that the participant but not the director could see their contents. The participant was asked to follow the instructions given by the director. For each trial, the director instructed the participant to move one of the eight objects by one slot (up, down, left, or right). Using a computer mouse, participants were required to click on the object they thought the director was referring to and to drag it into the appropriate slot on the shelves. Participants were given 3.6 seconds to make their response after each instruction.

There was one practice block followed by 16 test blocks, each made up of 3 trials. There were three types of trials. On filler trials, director’s instructions only referred to objects in non-occluded slots (i.e., visible to both director and participant). There were 2 filler trials per block and the third trial was either an experimental or a control trial. On experimental trials, the director instructed participants to move an object for which there were several types of the same kind on the shelves, with one of them in an occluded slot. For example, on one experimental trial, there were three balls of different sizes in different slots on the shelves, two of which were visible to the director. The director instructed participants to “move the small ball up.” The smallest ball of the three was in an occluded slot of the shelves, so the participant was required to consider the director’s perspective and move the smallest ball that the director could see, rather than the smallest ball that the participant themselves could see. Each experimental trial had a matched control trial in a different block wherein the objects on the shelves were identical to the experimental trial except that there was no longer an object of the same type in the occluded slot. The order of the 32 filler, 8 control and 8 experimental trials was counterbalanced between participants. A participant was removed from the analysis if they did not attempt a response on any experimental trials. Total scores were the mean accuracy across all eight experimental trials.

#### Attention and Reasoning Tasks

##### Attentional Network Test

The Attentional Network Test (Fan et al., 2002) was used to measure multiple aspects of attentional control. Each trial of the task began with an initial fixation period for 400-1600ms (black fixation cross centered on a grey screen), followed by a warning cue that was presented for 100ms. There were 4 possible kinds of warning cues: (1) an asterisk in place of the fixation cross, (2) two asterisks—one on the top and one on the bottom of the fixation cross, (3) one asterisk above the fixation cross, and (4) one asterisk below the fixation cross. The first two types of warning cues did not give information about the location of an upcoming target, whereas the latter two types of warning cues indicated the location of the upcoming target (above or below the fixation cross respectively). The target was a black arrow pointing left or right that appeared either above or below the fixation cross. The target was flanked on both sides by two arrows that pointed in the same direction (congruent condition), two arrows that pointed in the opposite direction (incongruent condition), or by two parallel lines (neutral condition). Participants were asked to indicate the direction of the target arrow by pressing one of two designated keys, using their left and right index fingers. Participants were asked to respond as quickly and accurately as possible to each trial. There were two blocks of 12 practice trials with feedback for accuracy. The first practice block did not have cueing, and the second practice block did have cueing. After the practice, participants completed 4 test blocks of 48 task trials comprising 12 trials of each cue condition.

Accuracy and reaction time (RT) data were collected from each block. Participants who scored lower than 50% accuracy on 2 or more blocks were excluded from analyses involving this task. Outlier detection for RTs was conducted within participants, such that trial RTs falling two standard deviations above or below the participant’s mean RT were excluded. Indices for attentional orienting and executive attention were calculated according to Fan et al., (2002), with the attentional orienting effect given by (mean RT on trials with a center cue) – (mean RT on trials with a spatial cue), and an executive attention effect given by (mean RT on trials with an incongruent flanker) – (mean RT on trials with a congruent flanker).

##### Matrix Reasoning Task

The color-blind friendly version of Matrix Reasoning Task (Chierchia et al., 2019) was used as a control measure of general reasoning ability. The task showed a 3 x 3 matrix on a screen the computer screen. Eight of the nine cells contained a colored abstract shape, and the bottom right cell of the matrix was empty. Participants were asked to complete the matrix by choosing one of 4 shape options displayed on the bottom of the screen. The shape characteristics varied in shape, color, size, and location. Each trial began with a 500ms fixation cross, followed by a 100ms white screen and participants had a time limit of 30s to complete each trial. A time warning was given 25s after the matrix was presented to indicate 5s remained in the trial. There were 3 easy practice trials followed by 8 minutes of test items presented in a fixed order. All items in practice and test had feedback on accuracy. There were 80 possible items in the whole set, but participants were not required nor expected to complete all items in the allotted time. If a participant completed 80 items before the testing time limit, the items were presented again in the same order until 8 minutes was reached, but responses for repeated items were not analyzed. The total Matrix Reasoning Task score was the number of correctly completed items per minute.

#### Self-Report Questionnaires

##### The Object-Spatial Imagery Questionnaire

The Object-Spatial Imagery Questionnaire (Blajenkova et al., 2006) is a 30-item questionnaire with 2 subscales that assess individual differences in visual imagery preferences and experiences for (1) objects and (2) spatial relations. Representing and processing spatial relations and transformations among objects are two important skills for spatial PT (Kozhevnikov & Hegarty, 2001). Thus, the present study administered the 15 items of the spatial relations subscale as a measure of self-reported spatial PT. The spatial relations sub-scale demonstrated good internal reliability previously (α =0.79; Blajenkova et al., 2006) and in the present study (α =0.79). Participants were asked to read each statement and rate how strongly they agreed or disagreed on a 5-point scale (“totally disagree” to “totally agree”). Total scores were obtained by averaging across the 15 items of the spatial subscale and therefore could range from 1 to 5. Higher scores indicated greater preference and experience for spatial-relational imagery.

##### Questionnaire of Cognitive and Affective Empathy

The Questionnaire of Cognitive and Affective Empathy (Reniers et al., 2011) is a 31-item questionnaire composed of items from four previously validated empathy questionnaires (Baron-Cohen & Wheelwright, 2004; Davis, 1983; Eysenck & Eysenck, 1978; Hogan, 1969). It has two subscales that measure cognitive empathy and three subscales that measure affective empathy. The authors defined cognitive empathy as comprehension of other people’s experience which requires representing, holding, and manipulating information in mind, whereas affective empathy is “the ability to vicariously experience the emotional experience of others” (Reniers et al., 2011). Thus, the two subscales that made up the cognitive empathy facet were used, as this is the construct of most relevance to the present study. The two subscales were perspective taking and online simulation, and they contained 10 and 9 items respectively. The Questionnaire of Cognitive and Affective Empathy cognitive empathy scale has demonstrated good internal reliability previously (perspective taking subscale: α = 0.85, online simulation subscale: α = 0.83, Reniers et al., 2011; total cognitive empathy scale: α = 0.85, Gomez et al., 2022) and in the present study (perspective taking subscale: α = 0.82, online simulation subscale: α = 0.84, total cognitive empathy scale: α = 0.86). Participants were asked to read each statement and indicate their agreement or disagreement on a 4-point scale (“strongly disagree” to “strongly agree”). Ratings for each item were summed for each subscale to get subscale scores. Total Questionnaire of Cognitive and Affective Empathy, cognitive empathy scores were found by summing the perspective taking and online simulation subscales. Thus, total scores could range from 19 to 76, with higher scores indicating greater self-reported affective PT.

##### The Mentalization Scale

The Mentalization Scale (Dimitrijević et al., 2018) is a 28-item questionnaire with 3 subscales: self-related mentalization, other-related mentalization, and motivation to mentalize. In the present study, the 10-item other-related mentalization subscale was used as a self-report indicator of cognitive PT. The other-related mentalization subscale demonstrated good internal reliability previously (α = 0.77; Dimitrijević et al., 2018) and in the present study (α = 0.70). Participants were asked to read each of 10 the statements and rate how correct each statement was about themselves on a 5-point scale (“completely incorrect” to “completely correct”). The total score of the Mentalization Scale other-related mentalization subscale could range from 10 to 50 and a higher score indicates better ability to mentalize about others.

## Results

Reported results reflect analyses conducted on accuracy of PT and reasoning tasks, summary scores from self-report measures, and reaction time indexes of attention. Descriptive statistics for all measures are listed in Table 2.

**Table 2.** Descriptive statistics for all behavioral and self-report measures.

| **Measure** | **n** | **Mean** | **SD** | **Min.** | **Max.** | **Skew** |
| --- | --- | --- | --- | --- | --- | --- |
| ANT - Orienting Index | 193 | 30.69 | 22.74 | -47.71 | 107.56 | 0.10 |
| ANT - Executing Index | 193 | 56.11 | 24.45 | -23.08 | 128.74 | -0.07 |
| MaRs – Number of Correct Items Per Minute | 193 | 3.29 | 0.70 | 1.75 | 5.25 | 0.34 |
| SOT - Average Angular Error | 193 | 43.18 | 25.18 | 6.67 | 113.08 | 0.52 |
| PTT-A - Number of Correct Items Per Minute | 193 | 3.95 | 1.44 | 0.67 | 9.00 | 0.41 |
| Strange Stories - Mental State Stories Score | 193 | 12.31 | 2.41 | 3.00 | 16.00 | -0.72 |
| Director Task - Mean Experimental Accuracy | 193 | 0.46 | 0.35 | 0.00 | 1.00 | 0.26 |
| EAT - Z-Scored Empathic Accuracy | 193 | 0.91 | 0.22 | 0.13 | 1.41 | -0.50 |
| Yoni - Number of Correct Affective Trials | 178 | 41.75 | 4.85 | 24.00 | 48.00 | -1.11 |
| OSIQ - Spatial Subscale | 193 | 2.48 | 0.58 | 1.07 | 3.93 | 0.15 |
| QCAE - Cognitive Empathy Subscale | 193 | 59.24 | 7.86 | 34.00 | 74.00 | -0.53 |
| MentS - Other-Related Mentalization Subscale | 193 | 40.39 | 4.02 | 26.00 | 50.00 | -0.49 |

*Note.* ANT = Attentional Network Test; MaRs = Matrix Reasoning; SOT = Spatial Orientation Test; PTT-A = Perspective Taking Task for Adults; EAT = Empathic Accuracy Task; OSIQ = Object Spatial Imagery Questionnaire; QCAE = Questionnaire of Cognitive and Affective Empathy; MentS = Mentalization Scale;

### Comparison of Performance on Control and Experimental Trials

For tasks that provided a control condition, performance was compared between experimental and control trials to check that typically expected behavioral patterns were observed with the present sample. Perspective Taking Task for Adults performance was assessed to determine if participants were quicker to respond to trials with 0° angular disparity (first-person perspective control trials) than those with 45°, 90°, 135°, and 180° angular disparities, including only trials for which a correct response was chosen. Paired samples t-tests indicated that mean reaction time for correct 0° angular disparity trials (*M* = 8.66s, SD= 3.24s) was significantly faster than mean reaction times for correct trials of each other angular disparity (*M*s > 10.8s; *p*s < .01; see Appendix, Figure A1 and Table A3).

Accuracy on Yoni physical trials was compared with accuracy on Yoni affective PT trials. Notably, fifteen participants scored at chance level or lower on Yoni physical accuracy trials but were not identified as outliers and did not fail any other comprehension checks in the experimental battery. These participants were not included in any analyses which involved Yoni Task performance but were retained in the dataset for inclusion in other analyses. Participants who passed the Yoni physical trials were significantly more accurate on physical (*M* = 0.92, SD = 0.11) than affective PT trials (*M* = 0.87, SD = 0.10) according to a paired-samples t-test, *t*(177) = 5.30, *p* < .001. Responses were also faster for physical (*M* = 2616.03ms, SD = 690.44ms) trials than for affective PT trials (*M* = 3728.51ms, SD = 981.45ms), *t*(177) = -22.22, *p* < .001.

Mean score of mental state stories and unlinked sentences trials were compared for the Strange Stories Task. A paired samples t-test indicated that participants scored significantly higher on unlinked sentences trials (*M* = 1.94, SD = 0.13) than on mental state stories trials (*M* = 1.54, SD = 0.30), *t*(192) = 17.23, *p* < .001.

To compare the effect of trial type (i.e., filler, control, and experimental) on accuracy for the Director Task, a one way within-subjects ANOVA test was used. There was a significant effect of trial type on accuracy, F(2, 220) = 330.68, *p* < 0.0001, η2(g) = 0.52. Post hoc pairwise comparisons indicated that all pairwise comparisons were significant. Mean accuracy for experimental trials (*M* = 0.46, SD = 0.35) was significantly lower than mean accuracy for filler trials (*M* = 0.95, SD = 0.04; t(192) = -19.4, p < .001) and mean accuracy for control trials (*M* = 0.90, SD = 0.10; *t*(192) = -17.4, *p* < .001). In addition, participants were significantly more accurate on filler than control trials, *t*(192) = 6.4, *p* < .001 (see also Appendix Figure A2).

Overall, comparison of performance on experimental and control trials for the tasks used in the present study aligned with expectations based on their original use (i.e., better performance or faster response times in control as opposed to experimental trials).

### Gender Differences in Performance

Group differences in performance by self-reported gender were investigated only for male and female participants, as there were too few non-binary participants (n=10) to include for comparison. Descriptive statistics for male and female participants are listed in Table 3. Mean differences for all measures and results of independent samples t-tests between genders are listed in Table 4. Males significantly outperformed females on the two behavioral and one self-report measure of spatial PT (*p*s <0.001), and the effect of gender was medium to large (see Table 4). Male participants also had significantly higher accuracy on the Director Task than female participants, with a small effect size (*d* = -0.33; see Table 4). Conversely, female participants had higher the Attentional Network Test orienting indices than males, again with a small effect size (*d* = 0.32). There were no significant gender differences on any other measures.

**Table 3.** Descriptive statistics by self-reported gender.

| Measure | Female |  |  |  |  | Male |  |  |  |  |
| --- | --- | --- | --- | --- | --- | --- | --- | --- | --- | --- |
|  | n | M | SD | Min. | Max. | n | M | SD | Min. | Max. |
| ANT - Orienting index | 120 | 33.23 | 22.97 | -30.52 | 107.56 | 63 | 25.97 | 21.82 | -47.71 | 74.11 |
| ANT - Executing index | 120 | 54.70 | 25.95 | -23.08 | 114.88 | 63 | 59.36 | 19.65 | 15.72 | 128.74 |
| MaRs – correct items per min. | 120 | 3.29 | 0.66 | 1.75 | 5.25 | 63 | 3.27 | 0.76 | 2.00 | 5.00 |
| SOT - average angular error | 120 | 50.22 | 23.88 | 7.42 | 108.33 | 63 | 30.84 | 23.44 | 6.67 | 113.08 |
| PTT-A - correct items per min. | 120 | 3.60 | 1.33 | 0.67 | 8.00 | 63 | 4.60 | 1.42 | 1.33 | 9.00 |
| Strange Stories - score | 120 | 12.25 | 2.45 | 3.00 | 16.00 | 63 | 12.32 | 2.38 | 6.00 | 16.00 |
| Director Task - accuracy | 120 | 0.42 | 0.32 | 0.00 | 1.00 | 63 | 0.53 | 0.38 | 0.00 | 1.00 |
| EAT - Empathic accuracy | 120 | 0.90 | 0.23 | 0.13 | 1.36 | 63 | 0.93 | 0.20 | 0.34 | 1.41 |
| Yoni - correct affective trials | 110 | 41.75 | 4.21 | 29.00 | 48.00 | 59 | 41.56 | 5.97 | 24.00 | 48.00 |
| OSIQ – Spatial | 120 | 2.30 | 0.53 | 1.07 | 3.93 | 63 | 2.86 | 0.51 | 1.80 | 3.87 |
| QCAE - Cognitive Empathy | 120 | 59.70 | 7.56 | 34.00 | 74.00 | 63 | 58.46 | 8.44 | 39.00 | 73.00 |
| MentS – Others | 120 | 40.62 | 3.79 | 29.00 | 49.00 | 63 | 39.79 | 4.31 | 26.00 | 49.00 |

*Note.* ANT = Attentional Network Test; MaRs = Matrix Reasoning; SOT = Spatial Orientation Test; PTT-A = Perspective Taking
Task for Adults; EAT = Empathic Accuracy Task; OSIQ = Object Spatial Imagery Questionnaire; QCAE = Questionnaire of
Cognitive and Affective Empathy; MentS = Mentalization Scale

**Table 4.** Group differences by self-reported gender.

| Measure | Mean difference | df | *p* | Cohen's *d* | Confidence Interval |
| --- | --- | --- | --- | --- | --- |
| ANT - Orienting Index | 7.26 | 181 | 0.04 | 0.32 | [0.02, 0.63] |
| ANT - Executing Index | -4.66 | 181 | 0.21 |  |  |
| MaRs – Correct Items Per Minute | 0.02 | 181 | 0.87 |  |  |
| SOT - Average Angular Error | 19.38 | 181 | <0.001 | 0.82 | [0.50, 1.14] |
| PTT-A - Correct Items Per Minute | -1.00 | 181 | <0.001 | -0.74 | [-1.05, -0.43] |
| Strange Stories - Score | -0.07 | 181 | 0.86 |  |  |
| Director Task - Accuracy | -0.11 | 181 | 0.04 | -0.33 | [-0.64, -0.02] |
| EAT - Empathic Accuracy | -0.03 | 181 | 0.25 |  |  |
| Yoni - Correct Affective Trials | 0.19 | 167 | 0.52 |  |  |
| OSIQ - Spatial | -0.56 | 181 | <0.001 | -1.09 | [-1.41, -0.76] |
| QCAE - Cognitive Empathy | 1.24 | 181 | 0.31 |  |  |
| MentS - Others | 0.83 | 181 | 0.18 |  |  |

*Note.* Degrees of freedom and *p*-values are reported for independent samples *t*-tests comparing performance of each measure between male and female participants. Significant differences are accompanied by Cohen's *d* effect size and a 95% confidence interval. Mean difference = mean female score – mean male score.

### Convergent Validity of Perspective Taking Within Domains

Convergent validity within perspective-taking domains was investigated by conducting Pearson’s correlations between each task and self-report measure within each domain (see Table 5). The two spatial PT tasks had a highly significant correlation with each other (*r* = -0.5, *p* < 0.001), and both were correlated with the Object-Spatial Imagery Questionnaire self-report measure of spatial PT (*p*s < .001). Cognitive PT tasks had a small significant correlation with each other (*r* = 0.2, *p* = 0.04), but neither correlated with the Mentalization Scale self-report subscale for other-related mentalization (*p*s < 0.05). Finally, the two affective PT tasks were not significantly correlated with each other, nor was either task correlated with the Questionnaire of Cognitive and Affective Empathy self-report of cognitive empathy (*ps* > 0.05).

**Table 5.** Results of Pearson’s correlations between all measures.

| Measure | 1 | 2 | 3 | 4 | 5 | 6 | 7 | 8 | 9 | 10 | 11 |
| --- | --- | --- | --- | --- | --- | --- | --- | --- | --- | --- | --- |
| 1 SOT (error) | --- |  |  |  |  |  |  |  |  |  |  |
| 2 PTT-A | -0.5** | --- |  |  |  |  |  |  |  |  |  |
| 3 OSIQ | -0.3** | 0.4** | --- |  |  |  |  |  |  |  |  |
| 4 Director | -0.1 | 0.1 | 0.3** | --- |  |  |  |  |  |  |  |
| 5 Stories | -0.1 | 0.0 | 0.0 | 0.2* | --- |  |  |  |  |  |  |
| 6 MentS | 0.0 | 0.0 | 0.1 | 0.1 | -0.1 | --- |  |  |  |  |  |
| 7 EAT | -0.3** | 0.1 | 0.1 | 0.0 | 0.0 | 0.0 | --- |  |  |  |  |
| 8 Yoni | -0.2* | 0.1 | 0.0 | 0.0 | 0.0 | 0.1 | 0.0 | --- |  |  |  |
| 9 QCAE | 0.0 | 0.0 | 0.2** | 0.0 | 0.0 | 0.7** | 0.0 | 0.1 | --- |  |  |
| 10 Orienting | 0.0 | -0.1 | 0.0 | 0.1 | 0.1 | 0.1 | 0.1 | 0.1 | 0.2* | --- |  |
| 11 Executing | 0.0 | 0.0 | 0.1 | 0.0 | -0.1 | 0.1 | 0.2* | 0.0 | 0.0 | -0.1 | --- |
| 12 MaRs | -0.1 | 0.3** | 0.1 | -0.1 | 0.1 | 0.0 | 0.1 | 0.1 | -0.1 | 0.0 | -0.1 |

*Note.* n = 178 for correlations with Yoni. n = 193 for all other correlations. ANT = Attentional Network Test; MaRs = Matrix
Reasoning; SOT = Spatial Orientation Test; PTT-A = Perspective Taking Task for Adults; EAT = Empathic Accuracy Task; OSIQ =
Object Spatial Imagery Questionnaire; QCAE = Questionnaire of Cognitive and Affective Empathy; MentS = Mentalization Scale;
\*\* $p < .001$ , \* $p < .05$ .

### Association of Perspective Taking Between Domains

Pearson’s correlations of cognitive PT (Strange Stories, mean mental state stories score; Director Task, mean accuracy on experimental trials) and affective PT (Empathic Accuracy Task, z-scored mean empathic accuracy; Yoni, number of correct affective trials) tasks resulted in no significant correlations between the two domains (see Table 5). However, self-report measures of cognitive PT (Mentalization Scale, other-related mentalization subscale) and affective PT (Questionnaire of Cognitive and Affective Empathy, cognitive empathy scale) were strongly correlated, even after removing the effect of the Matrix Reasoning Task (*r* = 0.7, *p* < 0.001). The Mentalization Scale other-related mentalization subscale and Questionnaire of Cognitive and Affective Empathy cognitive empathy scale were not correlated with cognitive nor affective PT tasks.

Pearson’s correlations of cognitive and spatial PT (Spatial Orientation Task, angular error; Perspective Taking Task for Adults, number of correct items per minute) behavioral tasks also indicated no significant associations (see Table 5). There was a small significant correlation between mean accuracy on the Director Task and self-reported spatial PT as assessed by the Object-Spatial Imagery Questionnaire spatial subscale (*r* = 0.3, *p* <.001). The correlation between the Director Task and Object-Spatial Imagery Questionnaire spatial subscale remained significant after removing the effect of the Matrix Reasoning Task (*r* = 0.25, *p* <.01). No significant correlations were found between reaction time on Perspective Taking Task for Adults and cognitive PT tasks (*ps* <.05).

Pearson’s correlations of affective and spatial PT tasks resulted in two small significant correlations between Spatial Orientation Task angular error and performance on the two affective PT tasks. The correlation between the Spatial Orientation Task and Empathic Accuracy Task z-scored mean empathic accuracy remained significant after controlling for the Matrix Reasoning Task (*r* = -0.27, *p* < 0.01), as did the correlation between Spatial Orientation Task and the number of correct affective trials on Yoni ( *r* = -0.18 *p* = .01; see Figure 2).

**Figure 2.**
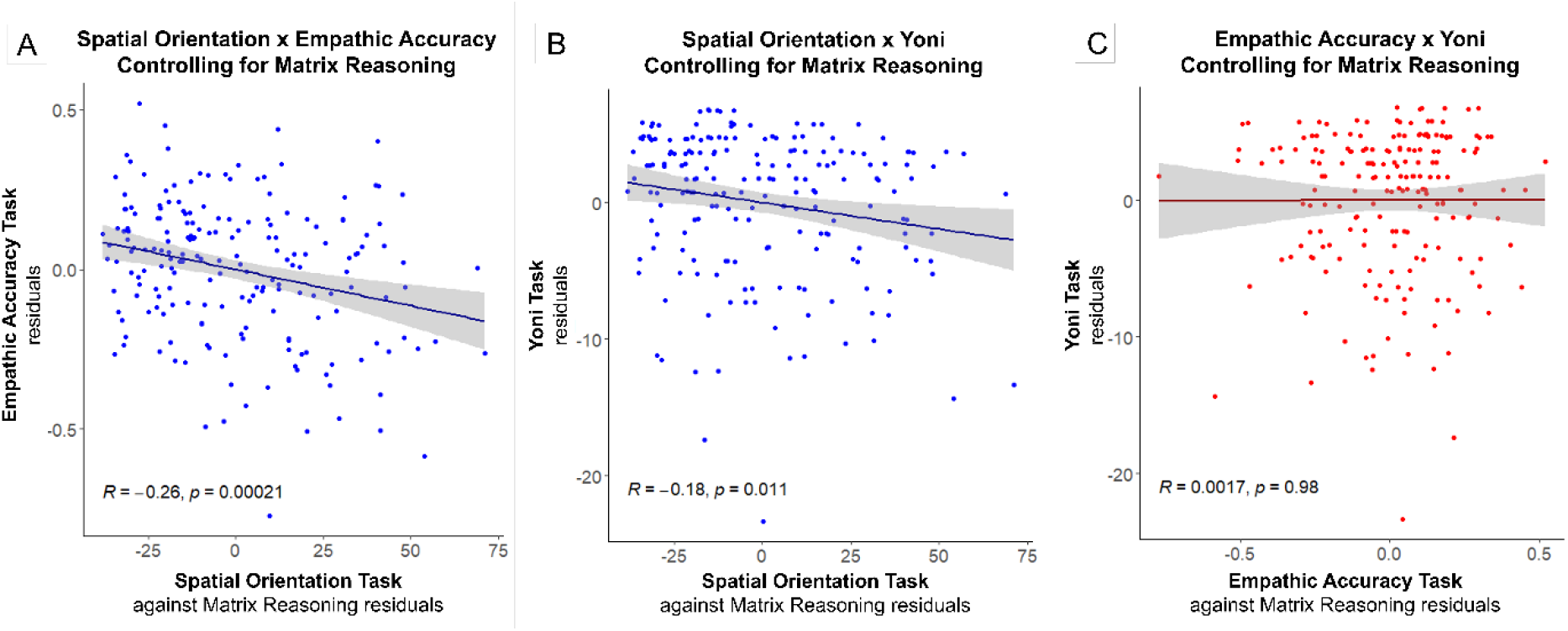
Scatter plots of the Spatial Orientation Task and affective perspective-taking tasks. *Note*. A) Includes data from 193 participants. B) Includes data from 178 participants. C) Includes data from 178 participants. All scatter plots show partial regression holding Matrix Reasoning Task performance constant.

The number of correct items per minute on the Perspective Taking Task for Adults was not significantly correlated with performance on the Yoni Task nor the Empathic Accuracy Task. The Questionnaire of Cognitive and Affective Empathy, cognitive empathy scale was modestly correlated with Object-Spatial Imagery Questionnaire spatial subscale, and the small correlation remained after controlling for the Matrix Reasoning Task (*r* = 0.19, *p* = 0.009). Further investigation indicated that the partial correlation of Object-Spatial Imagery Questionnaire spatial subscale with Questionnaire of Cognitive and Affective Empathy cognitive empathy scale, removing the effect of the Matrix Reasoning Task, was only significant for items in the Questionnaire of Cognitive and Affective Empathy perspective-taking subscale (*r* = 0.19, *p* = 0.01), but not for items in the Questionnaire of Cognitive and Affective Empathy online simulation subscale (*r* = 0.12, *p* = 0.09).

### Association of Perspective Taking with Attention

The Attentional Network Test executing index had a small significant correlation with the Empathic Accuracy Task z-scored mean empathic accuracy (*r* = 0.17, *p* <0.05). No other significant correlations were found between the Attentional Network Test executing index and any other PT measure. Attentional Network Test orienting index had a small significant correlation with the Questionnaire of Cognitive and Affective Empathy self-report measure of affective PT (*r* = 0.17, *p* <0.05). No other significant correlations were found between the Attentional Network Test orienting index and any other PT measures in the experiment. Attentional Network Test executing and orienting indices were not significantly correlated with each other, which is in alignment with original reports from this paradigm (Fan et al., 2002).

## Discussion

The fundamental aim of this study was to test the behavioral co-variance of spatial, cognitive, and affective PT, as a way to adjudicate between common and distinct mechanisms accounts. Pairwise correlations between PT measures of each domain, controlling for general reasoning ability, showed no significant correlations between spatial and cognitive PT tasks, or cognitive and affective PT tasks, and only a very small (although significant) correlation between one spatial PT task (Spatial Orientation Task) and the two affective PT tasks. However, the affective PT tasks were not correlated with each other. We also tested one version of a common-mechanism approach by investigating if attentional control or task stimulus features might explain behavioral co-variance among PT tasks. Attentional orienting was correlated with only one PT task (Empathic Accuracy Task) and executing was not significantly correlated with any PT tasks.

Overall, this pattern of results is most consonant with two of the distinct mechanisms accounts of PT abilities: the “distinct” account and the “spatial vs social” account. First, no significant correlations were found between spatial PT and cognitive PT tasks in the present study. This result stands in contrast to a recent study which found that reaction time on a novel spatial PT task was significantly correlated with Strange Stories Task performance (Putko & Złotogórska -Suwińska, 2019). One explanation for the apparent inconsistency of these findings relates to the validity of tasks used to measure spatial PT. Putko and Złotogórska-Suwińska (2019) devised a novel spatial PT task for use in their study. Despite its face validity as a measure of spatial PT, no psychometric analyses were provided to support its construct validity. Notably, the authors chose to use reaction times in place of accuracy in their correlation analyses due to ceiling effects in their sample. This may suggest that their measure lacked items of sufficient difficulty to capture a range of individual differences in spatial PT ability, and leaves to question what psychological construct their reaction time data is an indicator of, especially as response times to each question averaged less than one second. Conversely, the spatial PT tasks used in the present study have previously demonstrated wide ranges of item difficulty and high discriminability, high convergent validity with other spatial PT tasks, divergent validity from other closely related constructs like object rotation (Brucato et al., 2022), and additionally demonstrated convergent validity in the present study.

Second, self-reported affective PT was uncorrelated with spatial PT performance. Several previous studies have found that self-reported affective PT as measured by the Interpersonal Reactivity Index (Davis, 1983) or Empathy Quotient (Baron-Cohen & Wheelwright, 2004) were significantly associated with greater accuracy or quicker reaction times on spatial PT tasks (Erle & Topolinski, 2015; Gronholm et al., 2012; Mohr et al., 2010; Sulpizio et al., 2015; Thakkar & Park, 2010; Tomei et al., 2017). In contrast to previous studies, the Questionnaire of Cognitive and Affective Empathy cognitive empathy scale was not significantly associated with objective performance on either spatial PT task in the present study. This is particularly surprising because the Questionnaire of Cognitive and Affective Empathy cognitive empathy scale contains items from both the Interpersonal Reactivity Index and the Empathy Quotient. However, the Questionnaire of Cognitive and Affective Empathy was created by conducting a principal components analysis of items from four individually validated empathy questionnaires, and only the items which successfully loaded onto one of its five factors was retained (Reniers et al., 2011). As a result, only five out of fourteen original items measuring cognitive empathy on the Interpersonal Reactivity Index were considered valid items according to Reiners and colleagues (2011) analysis. Therefore, the results of this study suggest that when using a self-report measure of cognitive empathy which only included the most convergently valid items from previous questionnaires, self-reported affective PT is no longer associated with spatial PT.

Third, self-reported mentalizing was also uncorrelated with spatial PT performance. A handful of previous studies have found associations between self-reported autistic traits and “social ineptitude” scores as measured via the Autism-Spectrum Quotient and performance on spatial PT tasks that have agents in their stimuli (Brunyé et al., 2012; Clements-Stephens et al., 2013; Shelton et al., 2012) but not with spatial PT tasks without agents (Job et al., 2021; Clements-Stephens et al., 2013). In the present study, we aimed to directly test the association of spatial PT with self-reported cognitive PT via the Mentalization Scale other-related mentalization subscale (Dimitrijević et al., 2018; Richter et al., 2021) rather than indirectly via self-reported autistic symptomology alone. Given that the Mentalization Scale other-related mentalization subscale was not associated with the Perspective Taking Task for Adults or the Spatial Orientation Task, results demonstrate that objective spatial PT performance on a task with or without agents is uncorrelated with self-reported cognitive PT abilities.

Fourth, despite the high correlation between self-reported cognitive and affective PT measures used in the present study, no significant correlations were found between the cognitive and affective PT tasks. A brief review of behavioral correlations among various empathy and theory of mind tasks provided in the introduction reported inconsistent results regarding their behavioral associations. However, many of these correlational studies were conducted between cognitive PT tasks (e.g., Strange Stories Task) and emotion recognition or emotional empathy tasks, and therefore do not necessarily speak to their association with affective PT tasks. Future studies which directly compare behavioral performance on validated cognitive and affective PT tasks may be useful for understanding the association of these abilities.

The conclusion that spatial PT is distinct leaves open to question why co-deficits of each type of PT have been documented in an array of clinical populations (Baron-Cohen, 2000; Yirmiya et al., 1998; Song et al., 2019; Hamilton et al., 2009; Bora & Berk, 2016). One possibility is that co-deficits of spatial, cognitive, and affective PT in populations such as autism spectrum disorder and major depressive disorder arise from diverse rather than common cognitive impairments. Future work may benefit from focusing efforts on understanding what varied general cognitive processes may contribute to each type of PT ability to devise interventions for the improvement of spatial, cognitive, or affective PT abilities in clinical populations.

### Is There Any Support for Shared Cognitive Processes Among PT Domains?

A few findings from the present study may be interpreted as providing partial, though clearly very limited, support for shared cognitive processes among some PT tasks in different domains. In particular, small significant correlations were found among one spatial PT task (i.e., Spatial Orientation Task) and both affective PT tasks (i.e., Empathic Accuracy Task and Yoni), even after removing the effect of general reasoning ability, and even though the latter two tasks did not correlate with one another. This finding is somewhat unexpected given that performance on emotion recognition tasks has previous been found to be uncorrelated with spatial PT performance (Chiu & Yeh, 2018; Martin et al., 2019). This apparent lack of accord with earlier studies is not entirely problematic, however, since the tasks used in the present study were not pure emotion recognition tasks, but rather, are thought to additionally require mentalizing and differentiating between emotional mental states (Mackes et al., 2018; Shamay-Tsoory et al., 2007). Such qualities might have driven the observed relationship with the Spatial Orientation Task. However, neither affective PT measure was correlated with the alternative spatial PT task (i.e., Perspective Taking Task for Adults), despite its high convergent validity with the Spatial Orientation Task. The overall results provide little support for the idea that spatial PT and affective PT rely on shared processing resources.

One cognitive process that proponents of unitary accounts have argued could be shared among these tasks is attentional control (Sun & Wang, 2014). However, the present study found that the Attentional Network Test orienting and executing indices were not associated with spatial PT task performance. Therefore, individual differences in attentional control do not appear to account for the association of Spatial Orientation Task with either affective PT task. While the relationship between the Spatial Orientation Task and the Yoni Task is particularly puzzling, an alternative explanation for why smaller angular errors in perspective-taking on Spatial Orientation Task showed a modest negative correlation with accuracy in emotional intensity ratings of others on the Empathic Accuracy Task may be that both required magnitude judgments. It has previously been suggested that a generalized magnitude system serves as a basis for the development of accurate spatial judgements (Newcombe et al., 2015). Some researchers have also proposed that adults use a generalized magnitude system to organize representations of increasing emotional intensity (Holmes & Lourenco, 2011; though, see Pitt & Casasanto, 2018 for critiques). Thus, future behavioral work may investigate if accuracy of magnitude judgements more generally can explain the association of performance on these tasks.

A small positive correlation was also observed between accuracy on one cognitive PT task (i.e., Director Task) and self-reported preference/experience for spatial-relational imagery, as measured by the Object-Spatial Imagery Questionnaire spatial subscale. However, no significant associations were found between the Director Task and objective behavioral measures of spatial PT, and the Object-Spatial Imagery Questionnaire spatial subscale was not correlated with the other cognitive PT task (i.e., Strange Stories). We suspect that the small correlation between the Director Task and the Object-Spatial Imagery Questionnaire spatial subscale may be best attributed to the spatial features of Director Task stimuli, as compared to Strange Stories which had no visuospatial task demands.

Self-reported affective PT as measured by the Questionnaire of Cognitive and Affective Empathy perspective-taking subscale showed a weak significant correlation with the Object-Spatial Imagery Questionnaire spatial subscale. The creators of the Questionnaire of Cognitive and Affective Empathy describe their cognitive empathy perspective-taking subscale as measuring an individual’s propensity for “intuitively putting oneself in another person’s shoes to see things from his or her perspective” (Reniers et al., 2011). Importantly, the creators use the term “see” metaphorically in their description, as items on their perspective-taking subscale pertain to understanding the emotional perspective of others (e.g., “I can tell if someone is masking their true emotion”) and do not include phrases such as “see” or “look” that could be misinterpreted by participants as addressing spatial PT abilities. Therefore, it is unclear why the Questionnaire of Cognitive and Affective Empathy perspective-taking subscale and the Object-Spatial Imagery Questionnaire spatial subscale were correlated in our sample. Regardless, the weak correlation of Questionnaire of Cognitive and Affective Empathy cognitive empathy scale and Object-Spatial Imagery Questionnaire spatial subscale does not offer ample support for a common mechanisms accounts of PT when also considering that Questionnaire of Cognitive and Affective Empathy cognitive empathy scale was not significantly correlated with either behavioral affective PT measure.

### The Need for Convergent Validity Within Cognitive and Affective PT Domains

The measures of spatial PT used in this study have previously shown strong psychometric properties and convergent validity (e.g., Brucato et al., 2022; Blajenkova et al., 2006), and the coherence of measurement in this domain is corroborated in the present study. However, intentional investigations into convergent validity among measures within the cognitive PT and affective PT domains, such as those used in the present study, are sparse. In this data set, convergent validity was minimal among measures of cognitive PT, and poor among affective PT measures. Specifically, the two behavioral measures of cognitive PT, the Director Task and the Strange Stories Task, were weakly correlated after controlling for general reasoning abilities (consistent with Navarro, 2021), but neither cognitive PT task exhibited a significant relationship with the Mentalization Scale, other-related mentalization subscale self-report. Future research that carefully assesses the psychometric properties of existing measures of individual differences in cognitive and affective PT in neurotypical adults might help us to better understand what psychological constructs these tasks measure, and what general cognitive processes may additionally support performance.

### Conclusion

Despite weaknesses in convergent validity identified in the present study, the present findings do not support the “common” or “cold vs hot” accounts of PT. Further establishment of convergent validity within cognitive and affective PT domains needs to be addressed before conclusions can be drawn about the tenability of “spatial vs social” and “distinct” accounts of PT. Future work should establish construct validity for tasks of cognitive and affective PT before returning to the question of their association with spatial PT and one another.

## Appendix

**Figure A1.**
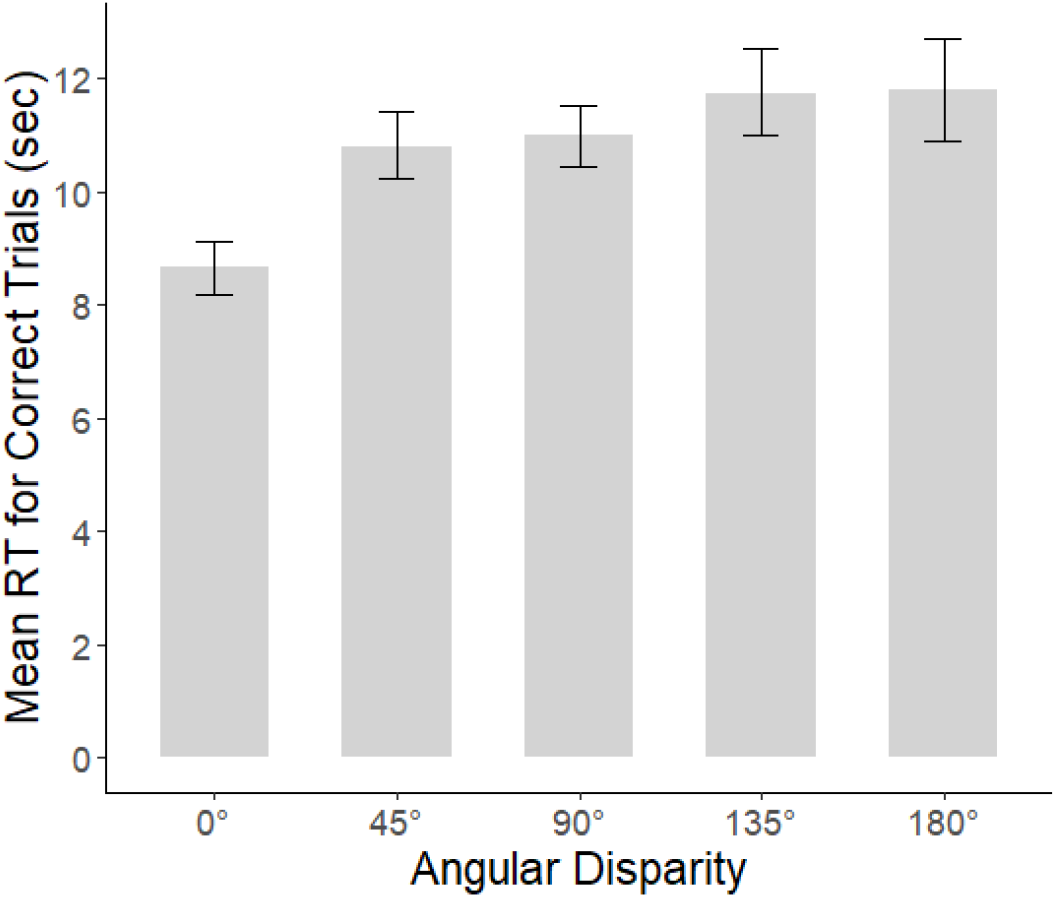
Mean RT by angular disparity on Perspective Taking Task for Adults trials. *Note*. RT = reaction time. Error bars represent ± 1 SEM.

**Figure A2.**
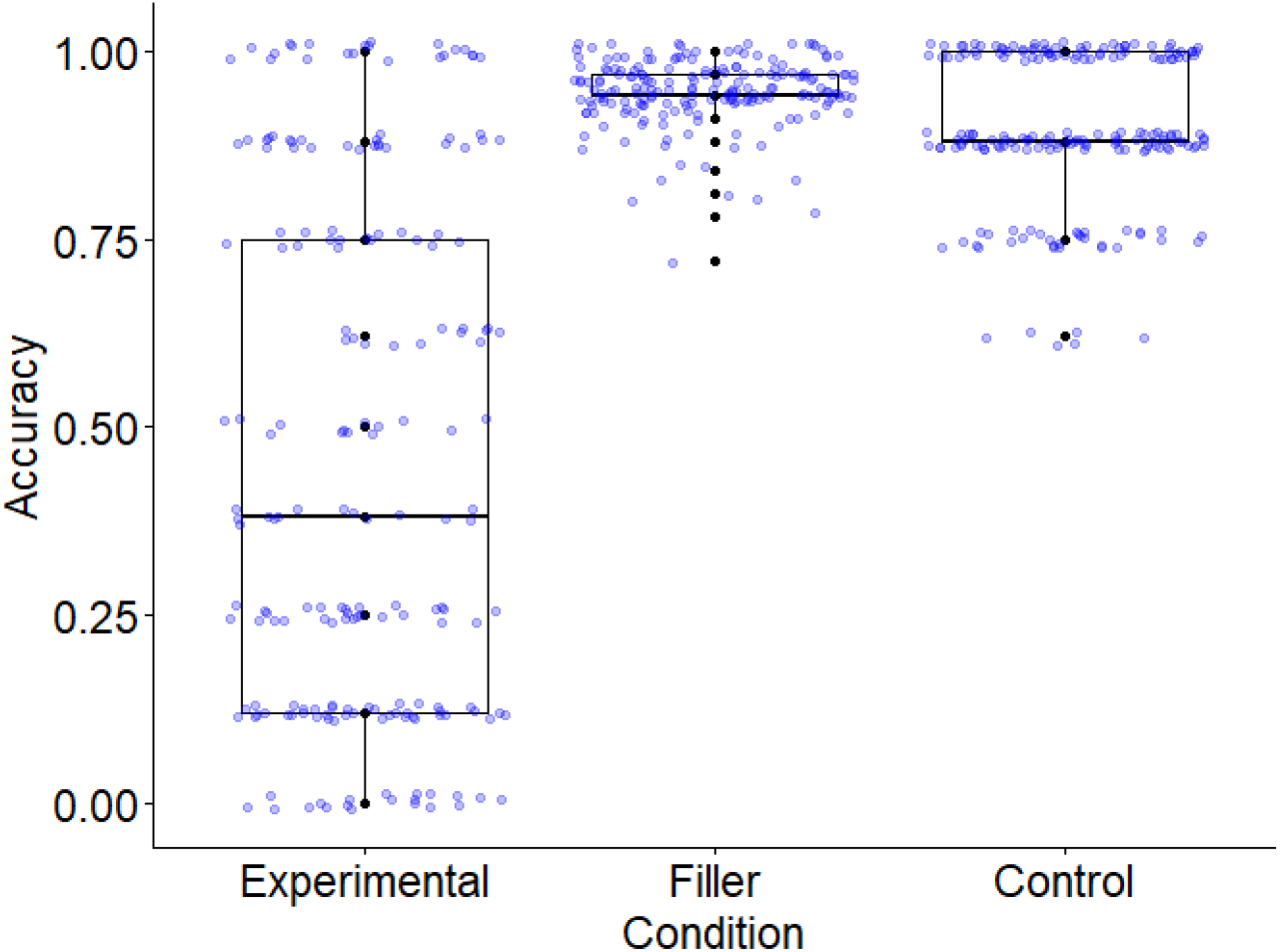
Mean accuracy by trial type on Director Task. *Note*. Blue dots are jittered data points of individual participants’ mean accuracy. Black dots are non-jittered data points. Experimental trials required the participant to consider the director’s perspective in order to achieve a correct answer. Control trials and filler trials did not require perspective taking for a correct response.

**Table A1.** Summary of behavioral studies with neurotypical adults on the association of spatial and affective PT.

| Year/<br>First Author | N | Spatial PT<br>Measure | Spatial PT<br>Measure:<br>Agent | Affective PT<br>Measure | Significant<br>Results | Control Task/<br>Condition |
| --- | --- | --- | --- | --- | --- | --- |
| Thakkar<br>2009 | 40 | OBT;<br>accuracy<br>& RT | Yes | IRI | Female ([RT back- RT front] x IRI,EC):<br>$r = .70, p = .0008$ | Line bisection x<br>IRI, $r = 0.55$,<br>$p = .0002$ |
| Thakkar<br>2010 | 32 | OBT;<br>accuracy<br>& RT | Yes | IRI | Female (accuracy front x IRI,PT):<br>$r = .63, p = .008$ | Letter rotation x<br>IRI, $r = .57, p = .03$ |
| Mohr 2010 | 100 | OBT;<br>accuracy<br>& RT | Yes | EQ (reverse-<br>scored) | Female (RT front x EQ):<br>$r = -.29, p = .04$ | NA |
| Gronholm<br>2012 | 85 | OBT;<br>accuracy<br>& RT | Yes | EQ (reverse-<br>scored) | PT Strategy Users ($n = 45$; RT x EQ,<br>total): $r = .413, p = .005$<br>PT Strategy Users ($n = 45$; RT x EQ,<br>ER): $r = .339, p = .023$ | NA |
| Erle 2015<br>(Exp. 1) | 113 | TFT;<br>accuracy | No | IRI | All (accuracy x IRI, PT) :<br>$r = .20, p = .033$ | NA |
| Erle 2015<br>(Exp. 2) | 120 | TFT;<br>accuracy | No | IRI | All (IRI, PT) :<br>$r = .20, p = .026$<br>All (accuracy x IRI, PT, removing effect<br>of Arithmetic): $r = .20, p = .031$ | Arithmetic ns<br>with IRI;<br>Arithmetic x<br>TFT, $r = .34, p < .001$ |

**Table A1. (continued)**
| Year/<br>First Author | N | Spatial PT<br>Measure | Spatial PT<br>Measure:<br>Agent | Affective PT<br>Measure | Significant<br>Results | Control Task/<br>Condition |
| --- | --- | --- | --- | --- | --- | --- |
| Erle 2015<br>(Exp. 3) | 169 | TFT;<br>accuracy | No | IRI | All (accuracy x IRI, PT):<br>$r = .22, p = .004$<br>PT Strategy Users (n = 86; accuracy x<br>IRI, PT): $r = .29, p = .008$ | NA |
| Erle 2015<br>(Exp. 4) | 163 | Table PT<br>Task<br>(Level 2);<br>RT | Yes | IRI | All (accuracy x IRI,PT):<br>$r = -.20, p = .013$ | NA |
| Sulpizio<br>2015 | 148 | Spatial<br>Memory<br>PT Task<br>(Level 2);<br>RT | Yes | IRI & BEES | All (non-avatar x BEES):<br>$r = -0.19, p < .05$<br>All (non-avatar x IRI,FS):<br>$r = -0.17, p < .05$<br>All (avatar 135° x BEES):<br>$r = -0.25, p < 0.01$ | Memory without<br>Spatial PT ns with<br>IRI & BEES |
| Tomei 2017 | 76 | Table PT<br>Task<br>(Level 2);<br># correct<br>per minute | Yes | IRI | All (correct/min x IRI,PT):<br>$\beta = 0.20, p = 0.04$<br>All (correct/min x IRI,PD):<br>$\beta = -0.23, p = 0.03$ | NA |
| Chiu 2018 | 158 | Spatial<br>Task-Switching;<br>switch<br>cost | No | RMET &<br>MET | All (switch cost x MET, emotional):<br>$\beta = -.32, p < .01$ | NA |

**Table A1. (continued)**
| Year/<br>First Author | N | Spatial PT<br>Measure | Spatial PT<br>Measure:<br>Agent | Affective PT<br>Measure | Significant<br>Results | Control Task/<br>Condition |
| --- | --- | --- | --- | --- | --- | --- |
| Martin 2019 | 146 | Street PT Task<br>(Level 1 &<br>Level 2);<br>accuracy & RT | Yes | RMET &<br>SEC | Younger Adults (allocentric CE<br>accuracy, Level 1 x SEC):<br>$r = -0.209$, $BF_{10} = 3.120$<br>All (allocentric CE accuracy, Level 2 x<br>SEC): $r = -0.244$, $BF_{10} = 9.904$ | Spatial PT with<br>traffic light ns with<br>SEC. |
| Nielsen<br>2015<br>(Exp.1) | 32 | Dot Task<br>(Level 1);<br>RT | Yes | IRI | All (avatar x IRI,PT):<br>$r = .32$, $p = .035$<br>All (avatar x IRI,EC):<br>$r = .34$, $p = .029$ | Spatial PT with<br>arrow ns with IRI. |
| Nielsen<br>2015<br>(Exp.2) | 28 | Dot Task<br>(Level 1);<br>gaze duration | Yes | IRI | All (avatar x IRI, PT):<br>$r = .37$, $p = .027$<br>All (avatar x IRI, EC):<br>$r = .32$, $p = .051$ | Dot Task without<br>avatar ns with IRI |
| Mattan<br>2016 | 43 | Dot Task<br>(Level 1);<br>accuracy & RT | Yes | IRI | All (Self-avatar CE RT x IRI, total):<br>$r = -.410$, $p = .008$<br>All (Self-avatar CE RT x IRI, PT):<br>$r = -.455$, $p = .003$<br>All (Self-avatar CE RT x IRI, EC):<br>$r = -.471$, $p = .002$ | NA |

*Note.* PT = perspective taking; OBT = Own Body Transformation Task; RT = reaction time; IRI = Interpersonal Reactivity Index; IRI, PT = perspective-taking subscale; IRI, FS = fantasy subscale; IRI, PD = personal distress subscale; IRI, EC = empathic concern subscale; EQ = Empathy Quotient; EQ, ER = emotional reactivity subscale; TFT = Tube Figures Test; BEES = Balanced Emotional Empathy Scale; RMET = Reading the Mind in the Eyes Task; MET = Multifaceted Empathy Test; SEC = Social-Emotional Cognition task; CE = Congruency Effect; ns = non-significant

**Table A2.** Summary of behavioral studies with neurotypical adults on the association of spatial and cognitive PT.

| Year/<br>First Author | N | Spatial PT<br>Measure | Spatial PT<br>Measure:<br>Agent | Cognitive PT<br>Measure | Significant<br>Results | Control Task/<br>Condition |
| --- | --- | --- | --- | --- | --- | --- |
| Shelton 2012<br>(Main exp.) | 72 | Three<br>Buildings<br>Task<br>(Level 2);<br>accuracy | Yes | AQ | All (agent x AQ, Social/Com.):<br>$r = -.584, p < .05$<br>Male (agent x AQ, Social/Com.):<br>$r = -.660, p < .05$<br>Female (agent x AQ, Social/Com.):<br>$r = -.501, p < .05$ | Object rotation, ns<br>with AQ;<br>Spatial PT without<br>agent, ns with AQ |
| Shelton 2012<br>(Replication) | 70 | Three<br>Mountains<br>Task<br>(Level 2);<br>accuracy | Yes | AQ | All (agent x AQ, Social/Com.):<br>$r = -.501, p < .05$ | NA |
| Brunyé 2012 | 140 | Table PT<br>Task (Level<br>1 & Level<br>2); RT | Yes | AQ | All (Level 2 angle effect x AQ, total): $\beta_{std} = .37, p < .01$<br>Female (Level 2 angle effect x AQ, total):<br>$\beta_{std} = .26, p < .05$<br>Male (Level 2 angle effect x AQ, total):<br>$\beta_{std} = .52, p < .01$ | NA |
| Clements-Stephens 2013<br>(Exp. 1) | 72 | Three<br>Buildings<br>Task<br>(Level 2);<br>accuracy | Yes | AQ; Social &<br>Com.<br>subscales | Agent trial presented first<br>(non-agent x AQ, Social/Com.):<br>$r = -.50, p = .001$<br>Agent trial presented first<br>(agent x AQ, Social/Com.):<br>$r = -.46, p = .004$<br>Agent trial presented second<br>(agent x AQ, Social/Com.):<br>$r = -.45, p = .008$ | When agent trial<br>presented second,<br>accuracy for non-agent trials ns with<br>AQ, Social/Com.. |

**Table A2. (continued)**
| Year/<br>First Author | N | Spatial PT<br>Measure | Spatial PT<br>Measure:<br>Agent | Cognitive PT<br>Measure | Significant<br>Results | Control Task/<br>Condition |
| --- | --- | --- | --- | --- | --- | --- |
| Clements-Stephens<br>2013<br>(Exp. 2) | 72 | Three<br>Buildings<br>Task<br>(Level 2);<br>accuracy | Yes | AQ; Social &<br>Com.<br>subscales | All (face agent x AQ, Social/Com.):<br>$r = -.40, p = .001$ | Spatial PT with<br>no-face agent ns<br>with AQ<br>Social/Com. |
| Clements-Stephens<br>2013<br>(Exp. 3) | 48 | Three<br>Buildings<br>Task<br>(Level 2);<br>accuracy | Yes | AQ; Social &<br>Com.<br>subscales | All (“triangle aliens” x AQ,<br>Social/Com.): $r = -.36, p = .013$ | NA |
| Job 2021 | 44 | Spatial<br>Orientation<br>Task (Level<br>2) Error | No | TSIS | NA | Object rotation ns<br>with TSIS. |
| Putko 2019 | 67 | Table PT<br>Task (Level<br>1 & Level<br>2);<br>accuracy &<br>RT | Yes | Strange<br>Stories Task | All (Level 2 RT x Strange Stories<br>accuracy): $r = -.36, p < .01$<br>All (Level 1 RT x Strange Stories<br>accuracy): $r = -.29, p < .05$ | Trail Making Task<br>and Wisconsin<br>Card Sorting Test<br>ns with spatial PT<br>or Strange Stories. |

*Note.* PT = perspective taking; AQ = Autism Quotient; AQ, Social/Com. = averaged social subscale and communications subscale; RT = reaction time; TSIS = Tromsø Social Intelligence Scale; ns = non-significant

**Table A3.** Descriptive statistics and paired t-tests for RT by angular disparity on PTT-A trials.

| Angular Disparity | n | Mean | SD | Min. | Max. | Comparison to 0° Trials |  |
| --- | --- | --- | --- | --- | --- | --- | --- |
|  |  |  |  |  |  | Mean Diff. | Conf. Int. |
| 0° | 191 | 8.66 | 3.24 | 3.77 | 21.09 | -- | -- |
| 45° | 188 | 10.80 | 4.14 | 4.29 | 34.95 | -2.20* | [-2.75 -1.65] |
| 90° | 191 | 10.99 | 3.93 | 5.62 | 30.09 | -2.24* | [-2.79 -1.70] |
| 135° | 189 | 11.74 | 5.34 | 4.44 | 41.76 | -3.11* | [-3.85 -2.37] |
| 180° | 183 | 11.82 | 6.19 | 3.98 | 42.06 | -3.29* | [-4.16 -2.41] |

*Note.* Descriptive statistics and paired samples t-test results with 95% confidence intervals comparing mean RT in seconds for the 0° angular disparity trials to the mean RT in seconds to each other trial type. RT = reaction time. PTT-A = Perspective Taking Task for Adults.
\* $p < 0.001$ .

